# Description of Life History and Reproductive Size Thresholds in Three High Elevation *Puya* (Bromeliaceae)

**DOI:** 10.1101/2022.02.18.480924

**Authors:** Leah Veldhuisen, Mary Carolina Garcia Lino, Erin Bodine, Julián Aguirre-Santoro, Rachel Jabaily

## Abstract

The Andes are a hotspot for biodiversity and high species endemism for both plants and animals. The genus *Puya* (Bromeliaceae) lives throughout the Andes, including puna and the páramo ecosystems above 3500m. Here, we studied the life history and reproductive size thresholds in three species of *Puya* (*P. raimondii, P. cryptantha* and *P. goudotiana*): *P. raimondii* in the Bolivian puna, and *P. cryptantha* and *P. goudotiana* in the Colombian páramo. We collected data on threshold size at flowering and clonal reproduction. All three species were found to have a consistent minimum size at flowering, while only *P. cryptantha* demonstrated a minimum size for clonal reproduction. No such evidence was found for *P. goudotiana*. We also found a positive correlation between leaf length and fruit number for *P. goudotiana* and *P. cryptantha*. Our data supported that *P. raimondii* is fully semelparous and indicated that *P. goudotiana* and *P. cryptantha* may be semi-semelparous.

## Introduction

The genus *Puya* Molina (Bromeliaceae) contains 228 species of rosette-forming terrestrial bromeliads endemic to Central and South America, from Costa Rica to Chile and Argentina (Gouda et al. 2018). *Puya* species are endemic to narrow regions at mid to high elevations from 1500 to 4000 m throughout the Andes, and are particularly species rich in high elevation páramos and inter-Andean valleys (Jabaily & Sytsma 2013). A phylogenetically distinct group of endemic species lives at low elevations down to sea level in central Chile (Jabaily & Sytsma 2010). *Puya* share a rosette body plan with the rest of the Bromeliaceae, and all produce terminal inflorescences with numerous, showy flowers (Mora et al. 2007, Smith & Young 1987). Species vary greatly in the size and leaf number at flowering time and overall size of floral display, which is particularly variable between sympatric *Puya* species in northern Andean páramos (Julian Aguirre-Santorro, personal communication). Species also differ in their ability - or inability - to produce clonal rosettes, a potential mode of asexual reproduction and the mechanism for continuation of the vegetative body after the inflorescence. Here, we characterize variation in these life history traits for three different *Puya* species, *P. raimondii* Harms, *P. goudotiana* Mez and *P. cryptantha* Cuatrec., in two different ecosystems.

Life history is often classified as semelparous (reproducing once during a lifetime) or iteroparous (reproducing multiple times), but this binary character may be more accurately described as a continuum of patterns of reproduction (Hughes 2017, Jabaily et al. 2021, Stearns 1992). *Puya* species span this continuum, and thus the genus may be a model for exploring life history evolution and adaptive life history features within the high Andean hotspots of biodiversity. In all species, the apical meristem produces the vegetative rosette, which switches from producing leaves and growing the vegetative body to making an inflorescence for sexual reproduction (Benzing et al. 2000). Each rosette produces one inflorescence; after the seeds are dispersed, the vegetative rosette stops growing and begins to senesce. The only mechanism for continuation of that genetic individual is for axillary meristems, located above the base of each leaf, to produce a new clonal ramet. Each ramet has its own apical meristem and potential to produce an inflorescence and additional ramets. The majority of *Puya* species produce many clonal ramets; individuals appear as a cluster of rosettes, potentially with many inflorescences.

Most *Puya* species are fully iteroparous, readily producing clonal ramets for many attempts at sexual reproduction. *Puya raimondii* is one of the few truly semelparous bromeliads, with individuals senescing without ever producing clonal ramets (Hornung-Leoni & Sosa 2005, Manzanares 2005). Some high-elevation *Puya* species also appear to have a decreased tendency to produce clonal ramets. Jabaily and Sytsma (2013) coined the term “semi-semelparous” to describe *Puya* species in which some individuals never produce clonal ramets prior to or after flowering and the individuals that do, only do so once they have transitioned to producing an inflorescence. Jabaily and Sytsma (2013) hypothesize that these species are in the process of evolving towards semelparity.

Both iteroparous and semelparous plants proceed through life cycle phases when they acquire and store enough energy to move forward, particularly for costly processes like sexual and asexual reproduction (Lacey 1986). Size is observed to be a strong predictor of the initiation of reproduction in a variety of species (Kuss et al. 2008, Lacey 1986, Miller et al. 2012, Mora et al. 2005, Werner 1975, Young 1984). For example, semelparous *Lobelia telekii* Schweinf. ex Engl., and iteroparous *Lobelia keniensis* Hemsl., two high elevation rosette plants on Mount Kenya, have a minimum size at sexual reproduction (Young 1990). Similarly, semi-semelparous *Puya hamata* L.B. Sm. has a uniform rosette diameter at the time of flowering (Garcia Meneses & Ramsay 2014).

In resource-limited habitats, the minimum size at reproduction will take longer to reach, and result in a longer life span, like that of long-lived semelparous plants (Young 1984, 1985). For many plants, both the likelihood of survival and flowering increase alongside plant size (Bonser & Aarssen 2009, Metcalf et al. 2003). This trend holds true in *Puya dasylirioides* Standl., with the number of mature fruits growing exponentially as rosette radius increases (Augspurger 1985). There is significantly less evidence for a size threshold in clonal or asexual reproduction, and evidence that does exist varies widely among genera and species (Ashmun & Pitelka 1985, Jabaily et al. 2021, Jabaily & Sytsma 2013, Mora et al. 2005, Schmid et al. 1995, Young 1984).

This research examines the life history traits of three *Puya* species: *P. raimondii, P. goudotiana* and *P. cryptantha. Puya raimondii* is the largest bromeliad in the world and is known as the ‘Queen of the Andes’, by far the largest plant found in the central Andean puna region, near 10 m when in flower (Dorst 1957). *Puya raimondii* is a dramatic example of semelparity; the species flowers once at 80-150 years old before senescing, and likely cannot produce clonal rosettes to continue the vegetative body (Lambe 2008). A *P. raimondii* inflorescence can be 4-6 m tall with potentially hundreds of thousands of yellow-ish flowers (Garcia Lino 2005, Hornung-Leoni & Sosa 2005, Sgorbati et al. 2003). It is endemic to the puna of central and southern Peru and northern and central Bolivia, generally above the elevation where other species of *Puya* in the region grow. The puna includes grass and shrubland above major forest belts between 3600 and 5000 m above sea level (Brush 1982, Morrone 2001). Populations of *P. raimondii* often live separately from each other in “rodales” on rocky hillsides in nutrient-poor soils (Augspurger 1985, Castillo et al. 2010, Garcia Lino 2005, Hornung-Leoni & Sosa 2005). The isolation of *Puya raimondii* populations has led to low genetic diversity, making it especially vulnerable to human and environmental threats (Lambe 2008, Sgorbati et al. 2003).

Further north in the Colombian Andes, the two other focal species of this study are endemic to the high-elevation páramo ecosystems: *P. cryptantha* and *P. goudotiana*. The páramo is similarly high in elevation (up to 4500 m), also above the forest belts, but has higher humidity and annual moisture levels with less seasonality (Balslev & Luteyn 1992, Luteyn 1999). The páramo is thought to be home to some of the fastest evolving lineages, with high species endemism (Madriñán et al. 2013). These two species grow in sympatry within the Cordillera Oriental of Colombia in multiple páramos, often with other *Puya* species (e.g. *P. nitida* Mez, *P. santosii* Cuatrec., etc.). They grow in boggy, water-logged areas and grasslands, sometimes at high regional population density. *Puya goudotiana* is one of the largest species of *Puya* after *P. raimondii*, reaching 5 m tall in flower with leaf lengths of over one meter (Smith & Downs 1974). Based on limited field observations, Jabaily and Sytsma (2013) categorized *P. goudotiana* as semi-semelparous along with other species from the páramos in Colombia and Ecuador. To our knowledge, no ecological studies have focused on *Puya goudotiana*. A 2005 study by Mora et al. in the Piedras Gordas section of Chingaza National Park found that *P. cryptantha* generally produces one to three clonal ramets at a leaf length between 8.8 and 20.8 cm, and usually flowers at a leaf length between 12.0 and 35.7 cm, making the species iteroparous.

Our goals in this study were to determine threshold leaf lengths for sexual and clonal reproduction, and to test for evidence of tradeoffs among vegetative body size and sexual and clonal reproduction in these three species. Physiological cues from body scaling for clonal and sexual reproduction in these specific species are unknown. For species that can reproduce clonally at some level like *P. cryptantha*, threshold size at reproduction is determined by a tradeoff between the benefits of beginning to sexually reproduce and the cost for future reproduction and survival. Semelparous species such as *P. raimondii* do not face this same tradeoff (Wesselingh et al. 1997). For clonal reproduction, there is only evidence of a minimum size in some species, and evidence of tradeoffs is unclear (Ashmun & Pitelka 1985, Jabaily et al. 2021, Mora et al. 2005, Schmid et al. 1995, Young 1984). We predict that all three species will have a minimum size at flowering, but *P. goudotiana* and *P. cryptantha* will not have a minimum size for clonal reproduction.

## Methods

### Study sites

To collect size and reproductive data on the three *Puya* species, we visited Bolivian puna and Colombian páramo habitats (Table 1, Figure 1). We collected data for *P. raimondii* in La Paz and Cochabamba, Bolivia in February 2018, and data on *P. cryptantha* and *P. goudotiana* in Chingaza National Park, near Bogotá, Colombia in October 2018.

**Figure 1.**
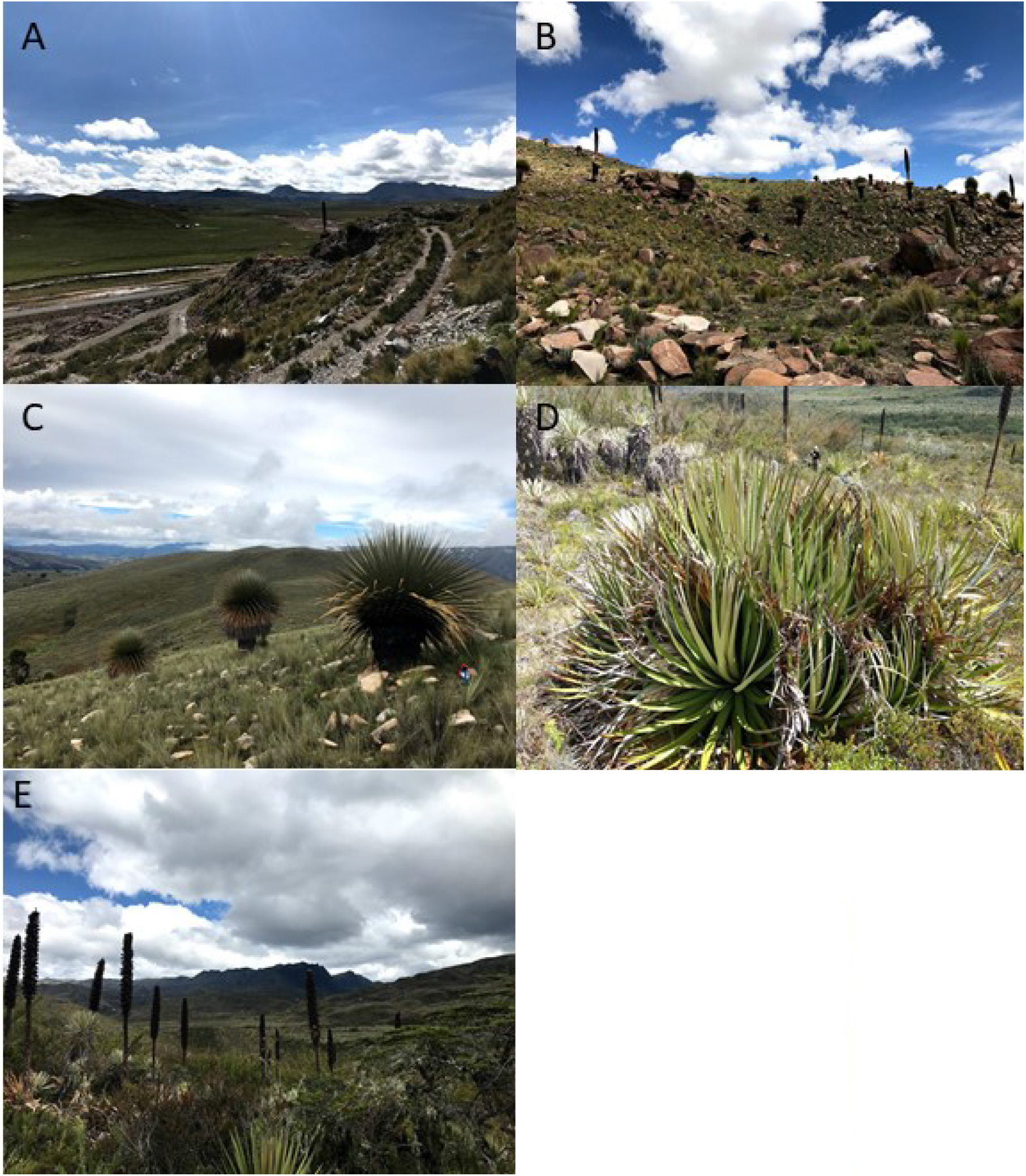
Photos **A-C** show *P. raimondii* in the puna of Bolivia. **D**. Many clonal ramets in *P. goudotiana* in the páramo. **E**. Post-flowering *P. goudotiana*. A-C, E by Leah Veldhuisen, D by Rachel Jabaily.

### Data collection

For *P. raimondii*, we constructed four plots of ten plants in each site using a randomly placed plot center, choosing general plot locations to cover the most space possible across the inhabited area. We thoroughly searched plots for small plants. For each individual, we measured four size traits: longest leaf length (LLL), longest leaf width (LLW), rosette height (RH), rosette width (RW), and noted the reproductive category (pre- and post-flowering) for the ten plants closest to the plot center. For each plot within a population, we also recorded elevation (see Table 1). For dead individuals, we only measured rosette height and reproductive category.

**TABLE 1.** Field measurement locations with GPS coordinates, elevation and species present.

| Site | *Puya* species present | Coordinates | Elevation (m) |
| --- | --- | --- | --- |
| **Comanche, La Paz, Bolivia** | *P. raimondii* | 16°96'09"S, 68°42'29"W | 4069 |
| **Totora Kasa, Cochabamba, Bolivia** | *P. raimondii* | 17°39'56.8"S, 65°35'09.4"W | 3822 |
| **Rodeo, Cochabamba, Bolivia** | *P. raimondii* | 17°23'44.5"S, 65°21'15.1"W | 3795 |
| **Toro Huarko, Cochabamba, Bolivia** | *P. raimondii* | 17°36'43.2"S, 65°32'18.9"W | 3307 |
| **Field Station, Cundinamarca, Colombia** | *P. goudotiana* | 4°31'32"N, 73°46'16"W | 3234 |
| **Chingaza Gate, Cundinamarca, Colombia** | *P. goudotiana*, *P. cryptantha* | 4°31'32"N, 73°42'24"W | 3383 |

Both leaf measurements were taken from the longest leaf, which we selected visually. We made all measurements using a 100-m transect tape. To measure the longest leaf length, we placed the tape on the leaf tip, and fed it into the center of the rosette. We measured the width one third of the way into the rosette on the same leaf. We measured rosette width with two people holding the transect tape alongside the plant. Rosette height was measured with a clinometer for the plants above 150 cm, and by holding the transect tape next to the plant for individuals below 150 cm. We determined reproductive category based on the presence of an inflorescence: pre- and post-flowering.

We collected data on size and reproductive category for *Puya goudotiana* and *P. cryptantha* using the same measurements. For *P. goudotiana*, we constructed two 5 by 5 m plots. We selected a random cardinal direction and a random number of steps to walk to determine the location of each plot. Within these plots, we measured all *P. goudotiana* individuals’ longest leaf length and width, rosette height and width, reproductive category and number of clonal ramets. We determined clonal ramets by proximity to larger rosettes. Ramet connections were not dug up, so we cannot definitively say they were clonal reproduction and not seedlings in close proximity. We excavated one with multiple attached rosettes to ensure that clonal reproduction was possible in this species.

Finally, we collected data on post-flowering *P. goudotiana* individuals. For plants with inflorescences, we measured longest leaf length and width, rosette height and width, inflorescence height, number of ramets, and estimates of the number of seeds capsules per inflorescence. The total number of capsules per inflorescence was estimated using capsule densities counted in 18 cm bands at the top, middle, and bottom of the flowering portion of the inflorescence. Assuming a linear increase in capsule density from the top to the middle, and a liner decrease in capsule density from the middle to the bottom, we were able to estimate the total number of capsules for an inflorescence as

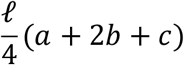

where *ℓ* is the total length of the flowering portion of the inflorescence, and *a, b*, and *c* are the capsule densities (measured in capsules/cm) at the top, middle, and bottom, respectively, of the flowering portion of the inflorescence. Figure 2 shows a graphical depiction of the estimated capsule density along the length of the flowering portion of the inflorescence. The formula for the total number of capsules is calculated as the total area under the curve from 0 cm to *ℓ* cm.

**Figure 2.**
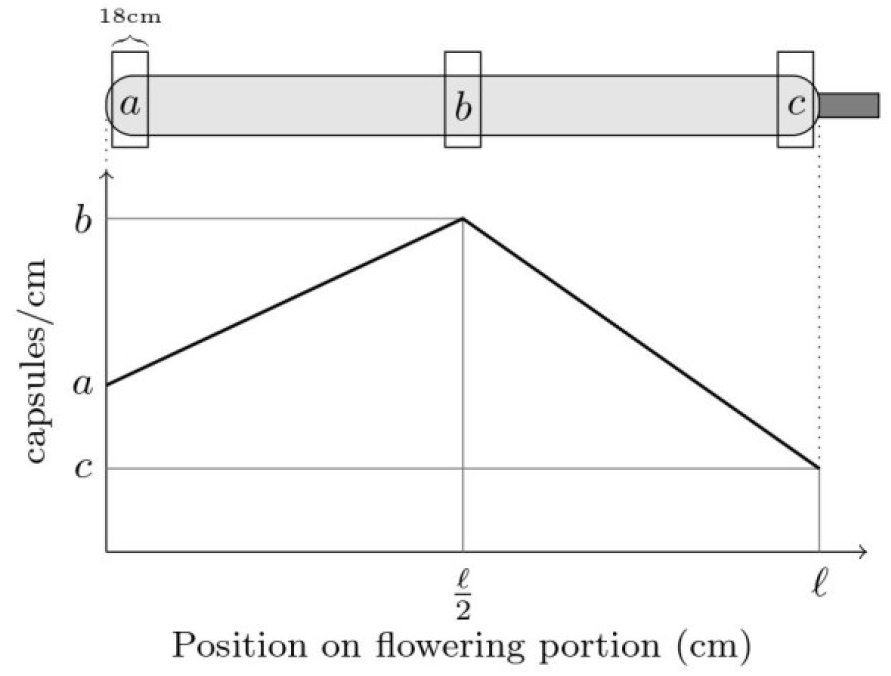
Graphical depiction of the estimated capsule density (measured in capsules/cm) based on capsule density measurements at the top, middle, and bottom of the flowering portion of the inflorescence.

We collected data on *P. cryptantha* for living individuals with inflorescences. We measured longest leaf length and width, total plant height, inflorescence circumference, number of pups, reproductive category, and number of fruits. Because the species was rare in the area we surveyed, and we only found post-flowering individuals that were submerged in a bog, we were not able to measure rosette height for any of them. We collected data on all plants in the area without constructing plots because there were few individuals.

### Analysis

We plotted all size metrics for the three species, and performed t-tests to check for differences between pre- and post-flowering individuals and those with and without ramets. We tested for a relationship between fruit number and leaf length using a linear regression. All analysis was performed in R v 4.1.1 (R Core Team 2021).

## Results

### Threshold size at flowering

All three species showed a threshold size for flowering (Figure 3). In *P. raimondii*, all size traits (LLL, LLW, RW and RH) were different for pre- and post-flowering individuals.

**Figure 3.**
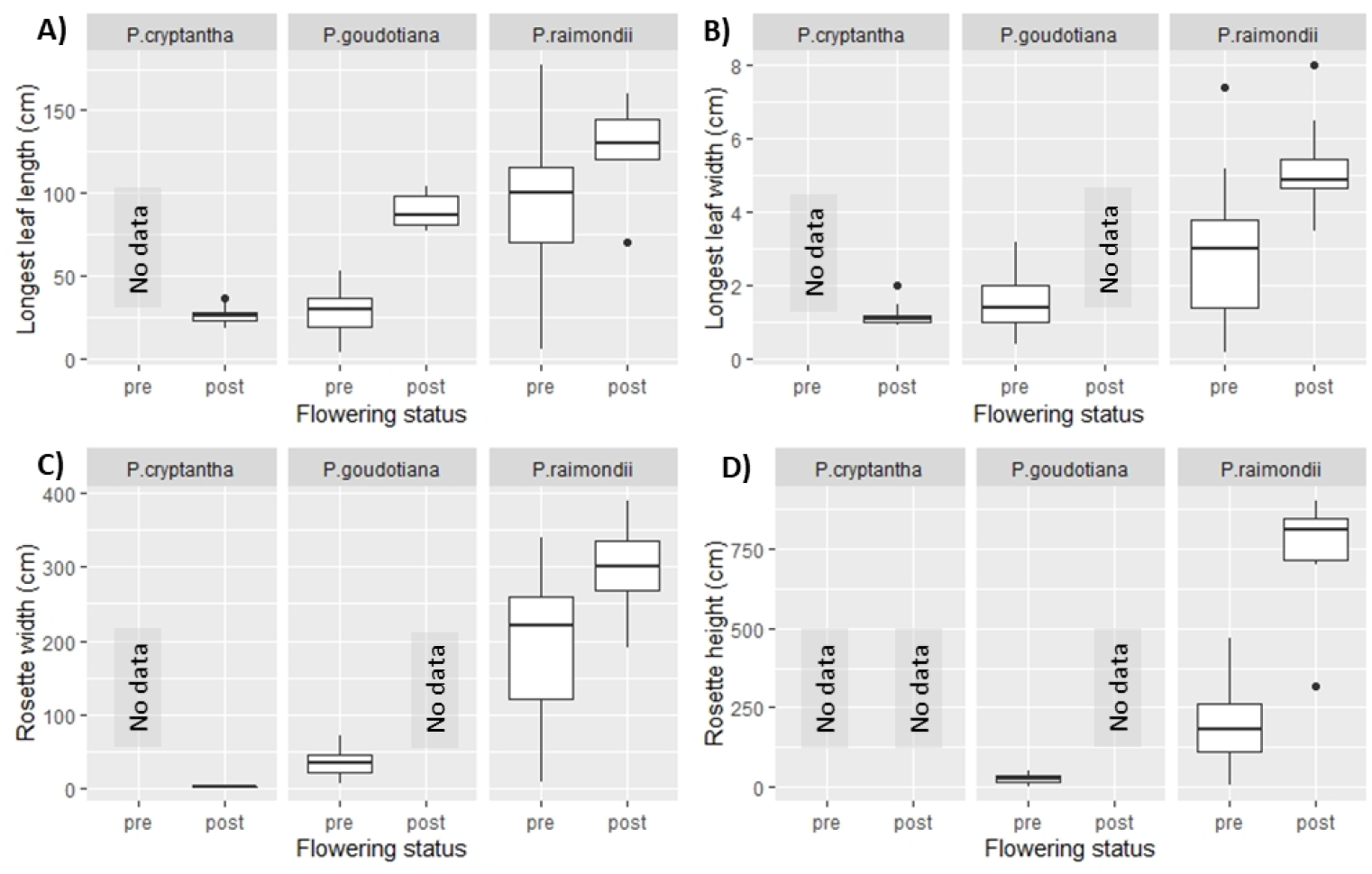
Boxplots showing each size trait for each species, divided by pre- and post-flowering individuals. **A**. Longest leaf length for all three species. **B**. Longest leaf width for all three species. **C**. Rosette width for all three species. **D**. Rosette height for all three species.

Longest leaf length and rosette height showed the most distinct difference between pre and post flowering (Figures 3A, 3D). All but one post-flowering individual had leaves longer than 120 cm; there were also many individuals that are above this size that had not yet flowered (Figure 3A; P<0.01). Similarly, all but one post-flowering individual had rosette heights above 650 cm (Figure 3D, P<0.01).

In *P. goudotiana*, longest leaf length (LLL) was the only trait measured in both pre and post flowering individuals. There was no overlap in this species in LLL between sizes of pre- and post-flowering individuals (Figure 3A; P<0.01). Only plants with leaves longer than 77 cm had flowered, and pre-flowering individuals did not have leaves longer than 53 cm.

Finally, *P. cryptantha* also seemed to show evidence of a threshold size for sexual reproduction, although we were not able to collect enough data to be certain. None of the flowering individuals had leaves less than 18 cm long, although all measured individuals had flowered (Figure 3A).

### Threshold for clonal reproduction

The two clonally reproducing species, *Puya cryptantha* and *P. goudotiana*, exhibited a less distinct minimum size for clonal reproduction than for flowering (Figure 4). For *P. cryptantha*, individuals with and without clonal ramets had leaves ranging from 18 to 36 cm, and there was no significant difference in leaf length between the groups (Figure 4A; P = 0.09). For the other two traits we measured in *P. cryptantha*, LLW and RW, there was also no difference in individuals with and without ramets (Figures 4B & 4C; P >0.28). In *P. goudotiana*, we also found no difference in any size trait between individuals with and without ramets (Figure 4A-D; P>0.1).

**Figure 4.**
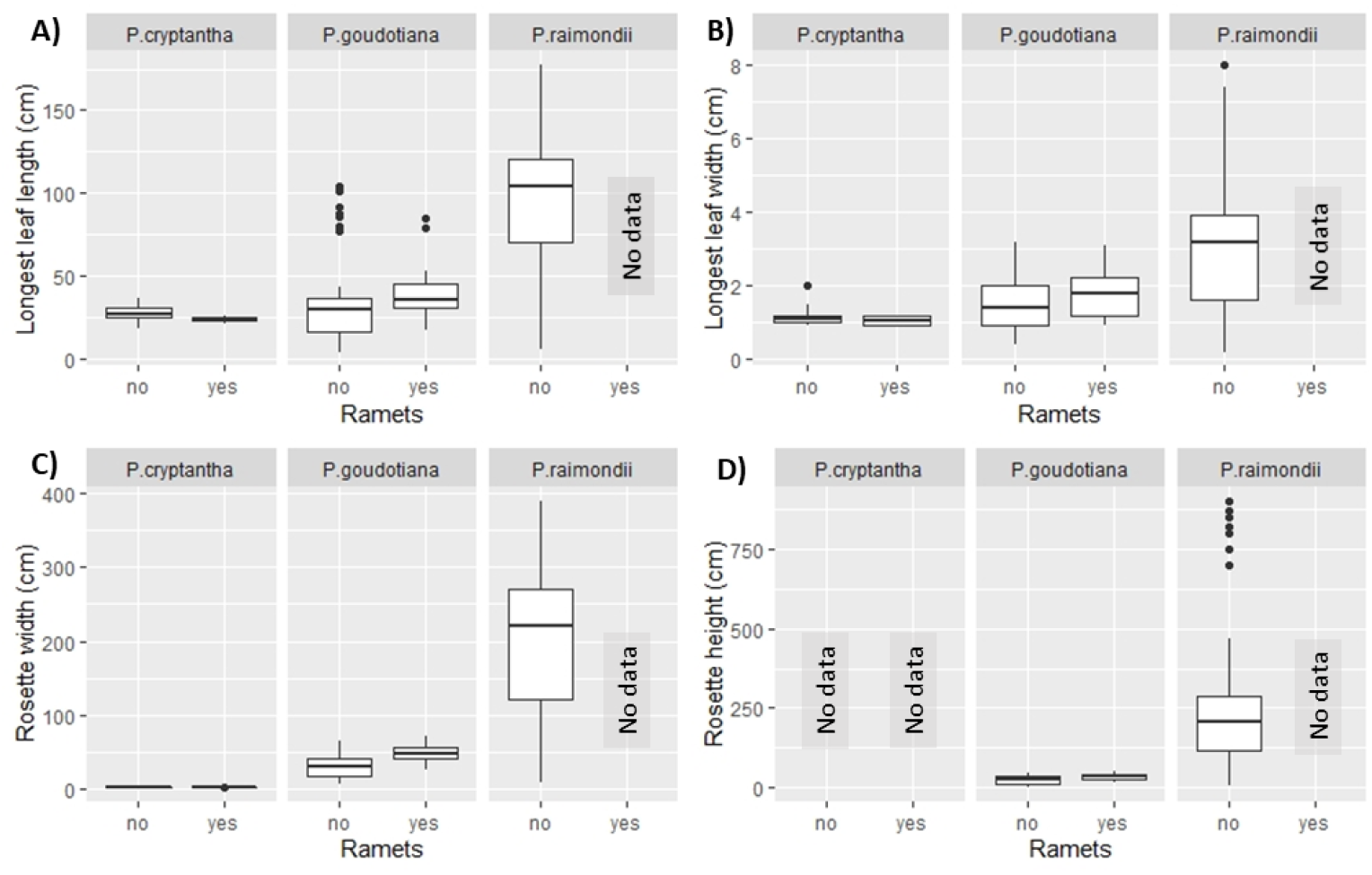
Boxplots showing each size trait for each species, divided by individuals with and without clonal ramets. **A**. Longest leaf length for all three species. **B**. Longest leaf width for all three species. **C**. Rosette width for all three species. **D**. Rosette height for all three species.

### Relationship between size and fruit number

Finally, we found a positive correlation between longest leaf length and number of fruits for *P. cryptantha* (Figure 5, R^2^=0.88).

**Figure 5.**
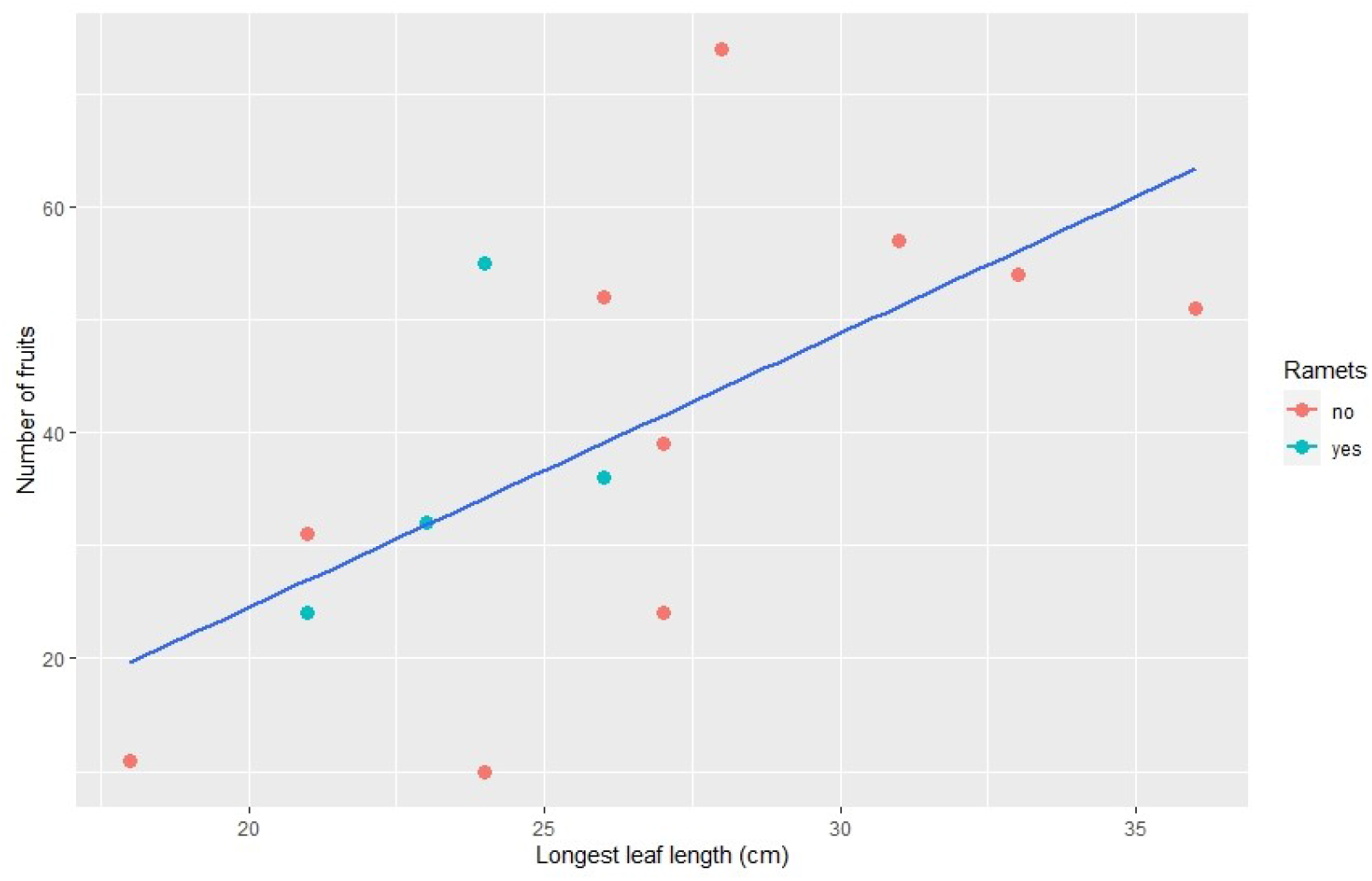
Number of fruits in *P. cryptantha* as predicted by longest leaf length (R^2^=0.88).

## Discussion

Overall, our results indicate a consistent narrow threshold vegetative size for sexual reproduction within all three species of *Puya*, as well as a threshold size for clonal reproduction in *P. cryptantha*. Additionally, we find evidence for a tradeoff in *P. cryptantha* between number of pups, leaf length and fruit number. Such a tradeoff is not visible in our data for *P. goudotiana*.

### Threshold for sexual reproduction for all species

The evidence for a threshold sexual reproductive size in the three studied *Puya* species differing in overall size, habitat, and life history, fits with a larger pattern of threshold size that has been described previously in other species (Augspurger 1985, Garcia Meneses & Ramsay 2014, Kuss et al. 2008, Mora et al. 2005, Young 1984). While a minimum size for flowering is an established phenomenon, reasoning for this threshold is less clear. This threshold may be due to meeting the minimum of stored fixed carbon, or the point when production of resources requires more energy than maintenance by the amount necessary to flower (Young 1984). Additionally, semelparous species’ inflorescence size is more dependent on rosette size at reproduction than iteroparous species, which would make minimum rosette size particularly important for *P. raimondii* (Young 1984). Multiple studies have found semelparous species’ reproductive output to be more sensitive to size at reproduction than that of iteroparous species (Schaffer & Schaffer 1977, 1979; Young 1990), but there is no evidence in our data that *P. raimondii* shows less variation in minimum flowering size than *P. cryptantha* or *P. goudotiana*. This may be because *P. raimondii* data were collected across a wider geographic range, or that *P. cryptantha* and *P. goudotiana* might be evolving towards semelparity.

Plants with larger flowers and less variable inflorescence size also tend to flower at larger sizes (Schmid et al. 1995), which may be the case for *P. raimondii* which has large, open flowers like other members of *Puya* subgenus *Puya*, in contrast to the typical narrow flowers of most other species of the genus. Semelparous *Yucca whipplei whipplei’*s reproductive output has been found to be highly responsive to increased photosynthate production from increased leaf surface area, while increased leaf surface area and photosynthate production had no impact on iteroparous *Yucca whipplei caespitosa*’s reproductive output (Huxman & Loik 1997). Although *P. cryptantha* and *P. goudotiana* are not entirely semelparous, minimum size at flowering is still significant and makes sense with prior studies in other genera (Mora et al. 2005, Young 1984).

### Threshold for clonal reproduction

Our data indicate a size threshold for clonal reproduction in *P. cryptantha*, similar to the findings of Mora et al. (2005), who more extensively studied the species. A minimum size for branching was determined in iteroparous *Lobelia keniensis* (Young 1984). Because *Lobelia* and *Puya* are comparable in their growth forms and tropical high elevation habitats, a clonal reproduction size threshold in *Lobelia* could suggest a similar, convergently evolved pattern in iteroparous *Puya* species.

### Tradeoffs to maximize reproductive output

Regardless of life history, all plants are partitioning their finite amount of energy towards competing goals of reproduction and individual growth, defense and maintenance. Our data show evidence of this tradeoff in *P. cryptantha*, as individuals with clonal ramets tend to be smaller and produce fewer fruits, while larger individuals have no clonal ramets but more fruits. For this tradeoff to benefit the plant, the clonal ramets must compensate with sufficient reproductive output to offset the loss from the shorter inflorescence of the mother rosette; larger inflorescences are strongly correlated with higher reproductive success, partially due to increased pollinators and a higher portion of set seeds (Huxman & Loik 1997, Inouye & Taylor 1980, Schaffer & Schaffer 1977, 1979; Young 1990). It is unknown if the clonal ramets of *P. goudotiana* or *P. cryptantha* flower independently of the mother rosette, which would be important to consider when analyzing this potential tradeoff. Iteroparous *Lobelia keniensis* inflorescence size does not increase with higher soil moisture, but the number of rosettes per individual does, indicating that the clonal ramets are of significant importance to the plant, and are the preferable investment (Young 1990).

### Life history categorization

*Puya raimondii* is documented as a semelparous species, as it never reproduces clonally, and dies after flowering (Smith & Downs 1974). Our data match this expectation, as *P. raimondii* individuals that we visited had one terminal inflorescence and did not have ramets, and all with inflorescences appeared dead or senescing. *Puya cryptantha* and *P. goudotiana* both reproduce via ramets and flowers, but their life history strategies are harder to categorize as semelparous or iteroparous. The term “semi-semelparity,” coined by Jabaily and Sytsma (2013), may be the correct categorization for both *P. cryptantha* and *P. goudotiana*, as both species have significantly reduced cloning ability compared to low-elevation *Puya* species and sympatric *P. nitida*. Some individuals of *P. goudoitana* and *P. cryptantha* were found flowering as a single rosette with no attached ramets. Additionally, we never saw either species’ clonal ramets flower on their own, indicating that they may have lost the ability to flower while evolving towards true semelparity. It is hard to tell if a flowering rosette is the initial seed-grown mother, or if the flowering rosette was the product of clonal reproduction from a long-decayed mother.

### Environment as a factor in life history evolution

While it has been established that *P. raimondii* is a semelparous species, there is currently no consensus of why this extreme life history strategy evolved in the species (Padilla 1973, Smith & Downs 1974). The evolutionary history of *P. cryptantha* and *P. goudotiana*’s transitional semi-semelparous life history strategies is similarly in need of further study. From other comparative studies, there is evidence of semelparity being associated with a drier site of one ecosystem, or the drier of two similar ecosystems (Young 1984). This mirrors the trend found in the three *Puya* species studied here, as the puna is significantly drier than the páramo and is the home of the only fully semelparous *Puya* species. Similarly, Young and Ausgsperger’s (1991) bet-hedging model for semelparity evolution suggests that highly variable and unpredictable environments may favor semelparity, especially in habitats prone to unpredictable drought like the puna. This trend offers a plausible explanation for why so many iteroparous and non-fully semelparous species inhabit the páramo, as its rainfall is consistent throughout the year and experiences significantly less seasonality than the puna. Based on Young’s 1990 study, plants with high survivorship, frequent reproduction and wet habitats require 13 times more reproductive output than iteroparous species to develop semelparity. Plants in dry sites with low survivorship and less frequent reproduction only require five times the reproductive output to evolve semelparity (Young 1990). This finding is supported by the fact that semelparous plants are more resource-dependent than their iteroparous counterparts (Huxman & Loik 1997; Young 1984, 1990). For semelparous plants, resources are essential to building an inflorescence, which takes longer in the resource-limited environments where large semelparous plants are common (Smith & Young 1987, Young 1990). Giant inflorescences are predicted by Young and Augsperger’s (1991) reproductive effort model of semelparity, which states that ever-increasing reproductive effort may favor semelparity, as larger inflorescences can be favored by pollinators and thus result in increased reproductive output (Rocha et al. 2005; Schaffer & Schaffer 1977, 1979). While pollinators have not been found to be major factors in the reproductive effort model (Young & Augspurger 1991), their presence and activity may be important in further determining how semelparity evolved in *P. raimondii*, and may be convergently evolving in the páramo lineages.

### Future study

While our data address important questions about the evolution and life history of *P. raimondii, P. cryptantha* and *P. goudotiana*, there are still many areas for continued investigation. Specific age-class survivorship data for *P. raimondii* would allow for evaluation of the demographic and bet-hedging models of semelparity evolution. Data on pollinator preference and reproductive effort, output and success would also help to evaluate the reproductive effort model and clarify habitat and pollinator role in life history evolution for the *Puya* genus. Comprehensive data on the environmental conditions of the puna and páramo would also be useful. Finally, reproductive output and success for semelparous and iteroparous species has been quantified in other genera, and would be interesting to analyze for *Puya* to test for evidence of the tradeoff between semelparity and iteroparity (Young 1990).

## Acknowledgments

We would like to thank the generous donors who made this project possible: the Keller Family Venture Grants and matching funds, the OBE Department funds, and the JH Enderson Research Award from Dr. Sharon Smith. Additionally, we thank Dr. Jim Ebersole and Dr. Mary Carolina Garcia Lino’s family, and everyone on Team Colombia and Team Bolivia who made the field research possible. Finally, thank you to Kate McGinn for helping with data analysis, and Katrina Dlugosch, the Dlugosch lab group and Jocelyn Navarro for manuscript feedback.

## Literature Cited

Ashmun, J. W., & Pitelka, L. F. 1985. Population Biology of Clintonia borealis: II. Survival and Growth of Transplanted Ramets in Different Environments. Journal of Ecology, 73(1): 185–198. https://doi.org/10.2307/2259777.

Augspurger, C. K. 1985. Demography and Life History Variation of Puya Dasylirioides, a Long-Lived Rosette in Tropical Subalpine Bogs. Oikos, 45(3): 341–352. https://doi.org/10.2307/3565569.

Balslev, H., & Luteyn, J. L. 1992. Paramo: An Andean Ecosystem under Human Influence. Academic Press Limited, University of Michigan.

Benzing, D. H., Bennett, B., Brown, G., Dimmitt, M., Luther, H., Ramirez, I., Terry, R., & Till, W. 2000. Bromeliaceae: Profile of an Adaptive Radiation. Cambridge University Press, Cambridge, United Kingdom.

Bonser, S. P., & Aarssen, L. W. 2009. Interpreting reproductive allometry: Individual strategies of allocation explain size-dependent reproduction in plant populations. Perspectives in Plant Ecology, Evolution and Systematics, 11(1): 31–40. https://doi.org/10.1016/j.ppees.2008.10.003.

Brush, S. B. 1982. The Natural and Human Environment of the Central Andes. Mountain Research and Development, 2(1): 19–38. https://doi.org/10.2307/3672931.

Castillo, J. S., Baldarrago, F. C. D., Poma, I., & Raimondo, F. M. 2010. Diagnostico del estado actual de conservación de Puya raimondii en Arequipa-Perú. Quaderni di Botanica ambientale e applicata, 21:83–91.

Dorst, J. 1957. The Puya Stands of the Peruvian High Plateaux as a Bird Habitat. Ibis, 99(4): 594–599. https://doi.org/10.1111/j.1474-919X.1957.tb03051.x.

Garcia Lino, M. C. 2005. Estado de conservacion de Puya raimondii Harms en el valle de Araca. Municipio Cairoma, La Paz—Bolivia. Thesis, Universidad Mayor de San Andres.

Garcia Meneses, P. M., & Ramsay, P. 2014. Puya hamata demography as an indicator of recent fire history in the páramo of El Ángel and Volcán Chiles, Ecuador-Colombia. Caldasia, 36(1): 53. http://dx.doi.org/10.15446/caldasia.v36n1.43891.

Gouda, E.. J., Butcher, D., & Gouda, C. S. 2018. Encyclopaedia of Bromeliads, Version 4. Utrecht University Botanic Gardens, Utrecht, Netherlands.

Hornung-Leoni, C., & Sosa, V. 2005. Morphological variation in Puya (Bromeliaceae): An allometric study. Plant Systematics and Evolution, 256(1–4): 35–53. https://doi.org/10.1007/s00606-005-0302-z.

Hughes, P. W. 2017. Between semelparity and iteroparity: Empirical evidence for a continuum of modes of parity. Ecology and Evolution, 7(20): 8232–8261. https://doi.org/10.1002/ece3.3341.

Huxman, T. E., & Loik, M. E. 1997. Reproductive patterns of two varities of Yucca whipplei (Liliaceae) with different life histories. International Journal of Plant Sciences, 158(6): 778.

Inouye, D. W., & Taylor, O. R. 1980. Variation in generation time in Frasera speciosa (Gentianaceae), a long-lived perennial monocarp. Oecologia, 47(2): 171–174. https://doi.org/10.1007/BF00346816.

Jabaily, R. S., Oberle, B., Fetterly, E. W., Heschel, M. S., Sidoti, B. J., & Bodine, E. N. 2021. Refining Iteroparity with Comparative Morphometric Data in Bromeliaceae. International Journal of Plant Sciences, 182(7): 577–590. https://doi.org/10.1086/715484.

Jabaily, R. S., & Sytsma, K. J. 2010. Phylogenetics of Puya (Bromeliaceae): Placement, Major Lineages, and Evolution of Chilean Species. American Journal of Botany, 97(2): 337–356. https://doi-org.ezproxy1.library.arizona.edu/10.3732/ajb.0900107.

Jabaily, R. S., & Sytsma, K. J. 2013. Historical biogeography and life-history evolution of Andean Puya (Bromeliaceae). Botanical Journal of the Linnean Society, 171(1): 201–224. https://doi.org/10.1111/j.1095-8339.2012.01307.x.

Kuss, P., Rees, M., Ægisdóttir, H. H., Ellner, S. P., & Stöcklin, J. 2008. Evolutionary demography of long-lived monocarpic perennials: A time-lagged integral projection model. Journal of Ecology, 96(4): 821–832. https://doi.org/10.1111/j.1365-2745.2008.01374.x.

Lacey, E. P. 1986. Onset of reproduction in plants: Size-versus age-dependency. Trends in Ecology & Evolution, 1(3): 72–75. https://doi.org/10.1016/0169-5347(86)90021-2.

Lambe, Antonio. 2008. IUCN Red List of Threatened Species: Puya raimondii. IUCN Red List of Threatened Species. Available from: https://www.iucnredlist.org/en (accessed 07/02/2022).

Luteyn, J. 1999. Páramo Ecosystem. Missouri Botanical Garden. Available from: http://www.mobot.org/MOBOT/research/paramo_ecosystem/introduction.shtml (accessed 01/17/2019).

Madriñán, S., Cortés, A. J., & Richardson, J. E. 2013. Páramo is the world’s fastest evolving and coolest biodiversity hotspot. Frontiers in Genetics, 4. https://doi.org/10.3389/fgene.2013.00192.

Manzanares, J. 2005. Jewels of the Jungle, Bromeliaceae of Ecuador, Part II Pitcairnioideae (Vol. 2). Imprenta Mariscal, Quito, Ecuador.

Metcalf, J. C., Rose, K. E., & Rees, M. 2003. Evolutionary demography of monocarpic perennials. Trends in Ecology & Evolution, 18(9): 471–480. https://doi.org/10.1016/S0169-5347(03)00162-9.

Miller, T. E. X., Williams, J. L., Jongejans, E., Brys, R., & Jacquemyn, H. 2012. Evolutionary demography of iteroparous plants: Incorporating non-lethal costs of reproduction into integral projection models. Proceedings. Biological Sciences, 279(1739): 2831–2840. https://doi.org/10.1098/rspb.2012.0326.

Mora, F., Chaparro, H. A., Bonilla, M. A., & Vargas, O. 2005. Rasgos de historia de vida de Puya cryptantha una bromelia monocárpica perenne. Universidad Nacional de Colombia Unibiblos, Bogotá, Colombia.

Mora F, Chaparro A, Vargas O, Bonilla A. 2007. Dinamica de la germancacion, latenica de semillas y reclutamiento de plantulas en Puya cryptantha y P. trianae, dos rosetas gigantes de los paramos Colombianos. Ecotropicos 1: 31–40.

Morrone, J. 2001. A formal definition of the biogeographic Paramo-Punan subregion and its provinces, based mainly on animal taxa. Revista del Museo Argentino de Ciencias Naturales 3(1): 1–11. http://dx.doi.org/10.22179/REVMACN.3.105.

Padilla, V. 1973. Bromeliads. Crown Publishers, New York.

RStudio Team. 2021. RStudio: Integrated Development Environment for R. RStudio, PBC, Boston, MA. URL: http://www.rstudio.com/.

Rocha, M., Valera, A., & Eguiarte, L. E. 2005. Reproductive ecology of five sympatric Agave littaea (Agavaceae) species in central Mexico. American Journal of Botany, 92(8): 1330–1341. https://doi-org.ezproxy1.library.arizona.edu/10.3732/ajb.92.8.1330.

Schaffer, W. M., & Schaffer, M. V. 1977. The adaptive significance of variations in reproductive habit in the Agavaceae. Evolutionary Ecology 60(5): 261–276. https://doi.org/10.1007/978-1-349-05226-4_22.

Schaffer, W. M., & Schaffer, M. V. 1979. The Adaptive Significance of Variations in Reproductive Habit in the Agavaceae II: Pollinator Foraging Behavior and Selection for Increased Reproductive Expenditure. Ecology, 60(5): 1051–1069. https://doi.org/10.2307/1936872.

Schmid, B., Bazzaz, F. A., & Weiner, J. 1995. Size dependency of sexual reproduction and of clonal growth in two perennial plants. Canadian Journal of Botany, 73(11): 1831–1837. https://doi.org/10.1139/b95-194.

Sgorbati, S., Labra, M., Grugni, E., Barcaccia, G., Galasso, G., Boni, U., Mucciarelli, M., Citterio, S., Iramátegui, A. B., Gonzales, L. V., & Scannerini, S. 2003. A Survey of Genetic Diversity and Reproductive Biology of Puya raimondii (Bromeliaceae), the Endangered Queen of the Andes. Plant Biology, 6(2): 222–230. https://doi.org/10.1055/s-2004-817802.

Smith, A., & Young, T. 1987. Tropical Alpine Plant Ecology. Annual Review of Ecology and Systematics, 18:137–158.

Smith, L. B., & Downs, R. J. 1974. Flora Neotropica: Pitcairnioideae (Bromeliaceae). Organization for Flora Neotropica by The New York Botanical Garden, New York, NY.

Stearns, S. C. 1992. The Evolution of Life Histories. Oxford University Press, Oxford, United Kingdom.

Werner, P. A. 1975. Predictions of fate from rosette size in teasel (Dipsacus fullonum L.). Oecologia, 20(3): 197–201. https://doi.org/10.1007/BF00347472.

Wesselingh, R. A., Klinkhamer, P. G. L., de Jong, T. J., & Boorman, L. A. 1997. Threshold Size for Flowering in Different Habitats: Effects of Size-Dependent Growth and Survival. Ecology, 78(7): 2118–2132. https://doi.org/10.2307/2265949.

Young, T. P. 1984. The Comparative Demography of Semelparous Lobelia Telekii and Iteroparous Lobelia Keniensis on Mount Kenya. Journal of Ecology, 72(2): 637–650. https://doi.org/10.2307/2260073.

Young, T. P. 1985. Lobelia telekii Herbivory, Mortality, and Size at Reproduction: Variation with Growth Rate. Ecology, 66(6): 1879–1883. https://doi.org/10.2307/2937383.

Young, T. P. 1990. Evolution of semelparity in Mount Kenya lobelias. Evolutionary Ecology, 4(2): 157–171. https://doi.org/10.1007/BF02270913.

Young, T. P., & Augspurger, C. K. 1991. Ecology and evolution of long-lived semelparous plants. Trends in Ecology & Evolution, 6(9): 285–289. https://doi.org/10.1016/0169-5347(91)90006-J.

